# Distinct populations suppress or escalate intake of cocaine paired with aversive quinine

**DOI:** 10.1101/2024.07.01.601599

**Authors:** Rosalie E. Powers, Peter A. Fogel, Jayson H. Reeves, Pamela Madrid, Travis M. Moschak

## Abstract

**Background:** Only a subset of individuals who encounter drugs of abuse become habitual users. Aversive subjective effects like coughing and unpleasant taste are predictors for continued use. While several preclinical studies have explored self-administration involving aversive cues, none have simultaneously introduced aversion with the initial drug self-administration. We aimed to develop a clinically relevant model by pairing intravenous cocaine with intraoral quinine self-administration from the outset and investigating whether repeated exposure to an aversive stimulus would alter its hedonic value under laboratory conditions.

**Methods:** Twenty-seven male and female Sprague Dawley rats self-administered intravenous/intraoral (cocaine/quinine) for 2 hr/day over 14 days. This was followed by a 1-day quinine-only extinction session, a 3-day return to self-administration, and an intraoral infusion session to assess quinine taste reactivity (TR).

**Results:** We identified three distinct groups. The first self-administered very little cocaine, while the second sharply escalated cocaine intake. Both groups had similar aversive TR to quinine, suggesting that the escalating group did not habituate to the aversive cue but pursued drug despite it. We also identified a third group with high initial intake that decreased over time. This decrease predicted high aversive TR, and we argue this group may represent individuals who “overindulge” on their first use and subsequently find self-administration to be aversive.

**Conclusions:** Our novel model mimics real-world variability in initial interactions with drugs of abuse and yields three distinct groups that differ in self-administration patterns and aversive cue valuation.

## Introduction

Identifying populations vulnerable to substance use disorder is an increasing focus for prevention strategies, as only a subset of individuals who encounter drugs of abuse become habitual users (Miech et al., 2023; SAMHSA, 2022). Importantly, the subjective effects associated with initial drug use such as coughing, unpleasant taste, euphoria, etc., may be predictors for or against continued use (Urbán, 2010). Furthermore, studies have suggested that the initial aversive effects associated with drugs of abuse weaken with time (Friedman et al., 1985; Hahn et al., 1990).

However, while numerous studies have explored self-administration involving aversive cues, to our knowledge, none have simultaneously introduced aversion with the initial administration of the addictive substance. Instead, these studies introduced the aversive stimuli as a punisher after reliable self-administration had been established (Seif et al., 2013; Torres et al., 2017; Lüscher et al., 2020). Here, we sought to develop a more clinically relevant model by pairing intravenous cocaine with intraoral quinine self-administration from the outset. We also sought to determine if repeated exposure to an aversive stimulus paired with an addictive substance would shift the hedonic value of that stimulus under laboratory conditions, as suggested by clinical studies (Friedman et al., 1985; Hahn et al., 1990). For this, we used a commonly used method developed by Grill & Norgren (1978) to assess aversive and appetitive taste reactivity (TR) to an aversive quinine cue paired with cocaine self-administration.

## Methods

### Subjects

Subjects were twenty-seven Sprague Dawley rats (male=13, female=14) (Envigo, Indianapolis, IN) weighing 193-447g. Twenty-one rats had previous experience pressing a lever for sucrose pellets in contextually distinct chambers from the chambers used in this experiment. These rats showed no behavioral differences from experimentally naive animals (n=6) and their data were analyzed together. Rats were individually housed with a 12-hour light/dark cycle (lights on at 8pm). All experiments were conducted during the dark cycle. Rats had unlimited access to water and rat chow (Teklad Irradiated Global 18% Protein Rodent Diet, Inotiv, Madison, WI). Procedures were conducted in accordance with the National Institutes of Health Guidelines for the Care and Use of Laboratory Animals and in accordance with the University of Texas at El Paso Institutional Animal Care and Use Committee.

### Surgery

The experimental timeline is shown in Fig. 1A. Rats were first anesthetized with ketamine/xylazine (100 mg/kg ketamine/10 mg/kg xylazine i.p.). Then, a chronic indwelling catheter was inserted into the right jugular vein (Access Technologies, Skokie, IL, USA) with an access port affixed to the dorsal region subcutaneously. Simultaneously, an oral cannula consisting of PE 100 (ID: 0.86mm, OD: 1.52mm) polyethylene tubing (Thomas Scientific, Swedesboro, NJ, USA) was surgically implanted anterolateral to the first maxillary molars then secured into place using two Teflon washers (Sterling Seal & Supply, Neptune, NJ, USA). Rats had at least 7 days of recovery before starting the experiment.

**Figure 1.**
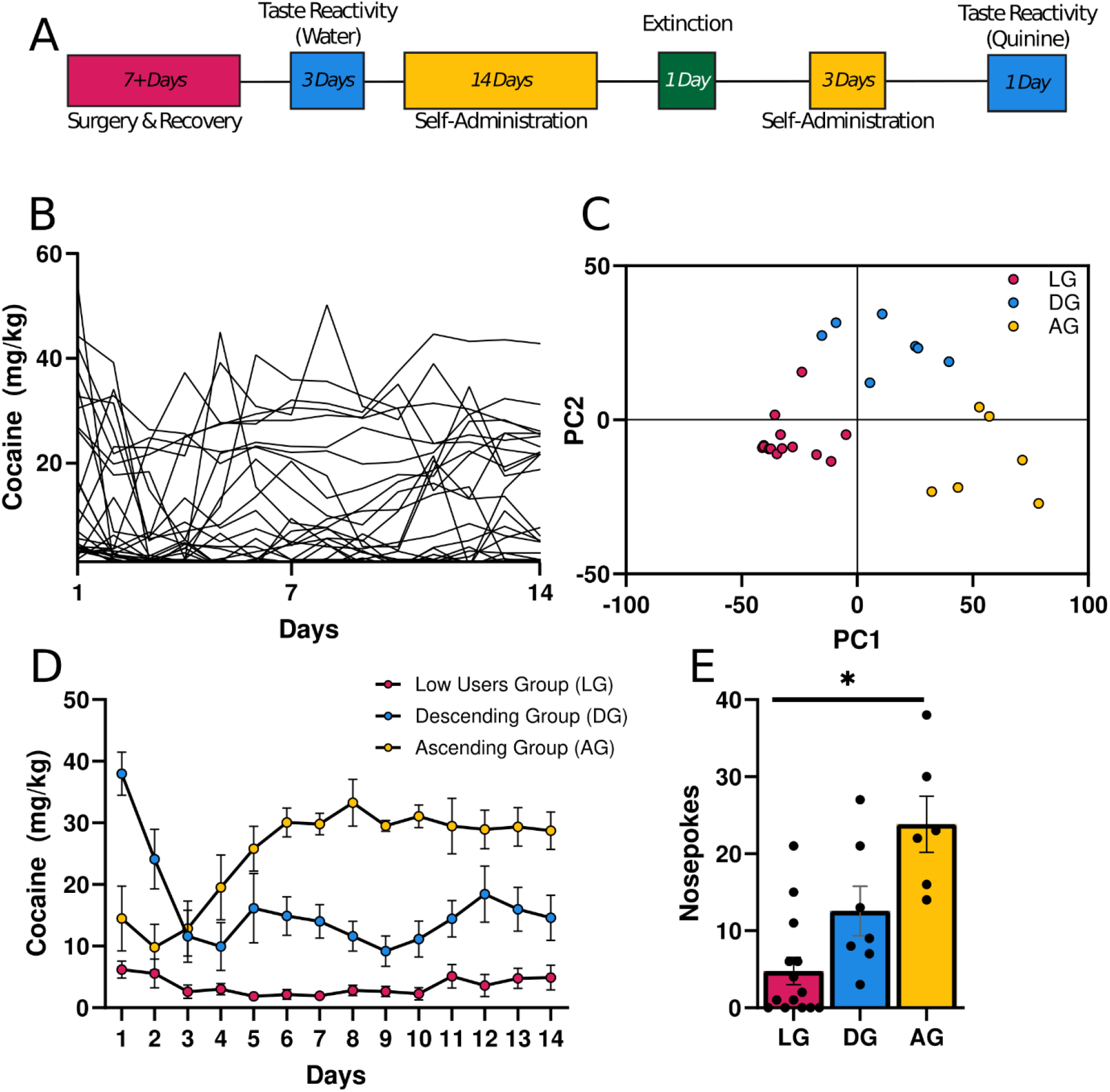
Distinct groups emerge during self-administration of a cocaine-quinine pair. **A)** Timeline of experimental procedures. Days for each task are included. **B)** Self-Administration. Amount of cocaine (mg/kg) infused during 14 days of self-administration for all rats (n=27). **C)** Principal Component Analysis of self-administration data. Principal Component 1 (PC1) is plotted against Principal Component 2 (PC2). Following k-means clustering, three distinct groups emerged: Low User Group (LG), Descending Group (DG), and Ascending Group (AG). **D)** Self-Administration. Average amount of cocaine (mg/kg) infused during self-administration of the LG, DG, and AG groups. As their names suggest, these groups either increased intake, decreased intake, or did not intake cocaine. **E)** Extinction. Similar to self-administration, the AG rats had the highest cocaine seeking behavior, followed by the DG rats and LG rats. * p < 0.05.

### Apparatus

Intraoral habituation and recording sessions were performed in a 44×43×72 cm behavioral chamber, whereas self-administration and extinction were performed in a smaller 29.5×25.5×31 cm chamber (Med Associates, St. Albans, VT, USA). Both chambers were housed in noise-reducing cubicles (Med Associates). Fluid delivery was accomplished through syringe pumps (Med Associates, St. Albans, VT, USA) with Tygon ND-100-80 tubing. Med-PC V software recorded and regulated all self-administration events and controlled intraoral fluid delivery. For TR, rats were placed on a clear platform situated 40 cm from the base of the chamber with an angled mirror underneath to allow visualization and recording.

### Habituation

Following recovery, rats underwent daily habituation sessions. Each session consisted of 45 intraoral water infusions on a VT-45 s schedule of delivery over approximately 40 minutes. Habituation ended when the rat gaped on less than 7 trials per session (max 5 days).

### Self-Administration

Self-administration consisted of 17 sessions separated into 2 discrete phases, as described in our experimental timeline. Each session was 2 hr and nosepokes resulted in i.v. cocaine hydrochloride (0.33mg/infusion) and intraoral quinine hydrochloride dihydrate (0.3mM, 0.13mL/infusion). Rats were randomly assigned left (n=14) or right (n=13) nosepokes.

### Extinction

Following 14 days of cocaine self-administration, rats underwent an extinction session. Extinction mimicked self-administration, except a nosepoke did not result in cocaine but instead only delivered intraoral quinine.

### Taste Reactivity Test

Following extinction, rats self-administered for three more days. Intraoral quinine followed the second self-administration phase. During the intraoral quinine session, rats received 45 intraoral quinine infusions on a VT-45 s schedule (0.3mM, 0.13mL/infusion). The session was recorded with a GoPro to quantify TR. One rat lost patency before completing intraoral quinine and was removed from the experimental protocol. Video recordings of intraoral quinine were scored by experimenters blind to experimental conditions. Rat behavior throughout infusions was analyzed by established criteria (Grill et al., 1978). Briefly, orofacial expressions following intraoral infusion were recorded as “gapes” (aversive) if they matched a “triangle” shape, and lateral tongue protrusions were recorded as “licks” (appetitive).

### Data Analysis

Principal component analysis (PCA) and k-means clustering were run on self-administration data to outline distinct groups (Fig. 1B,C). Cocaine intake was analyzed using a 14 × 2 × 3 (Day x Sex x Group) repeated-measures ANOVA. Separate univariate ANOVAs were run on extinction and TR data. Further analysis examined “net aversiveness,” defined as (number of gapes) + (number of licks * -1). Significant main effects or interactions were followed by Bonferroni corrected t-tests. For repeated-measures analyses that did not pass Mauchly’s test for sphericity, Huynh–Feldt-adjusted degrees of freedom were used. Pearson correlations were used to investigate the relationship between different behaviors. One set of correlations examined the relationship between the last 3 days of self-administration and TR. To investigate how the change in self-administration across days may have interacted with TR, another set of correlations examined the slope of the self-administration line for each rat and correlated it with TR. Grubbs’ test was used to check for significant outliers in all datasets. Datasets containing significant outliers underwent winsorization before further analysis.

## Results

### Self-Administration and Extinction

Cocaine intake was recorded for each animal (*Figure 1B*). PCA and k-means clustering were run on these data to define distinct groups (Fig. 1C). Daily cocaine for each group was plotted in Figure 1D. Based on their pattern of responding, these groups were labeled “Low Users Group” (LG), “Descending Group” (DG), and “Ascending Group” (AG). There was a significant interaction between group and day seen during self-administration (F(16.733,175.702) = 7.132, p<0.001). Individually, the LG group had no significant day effect (F(4.635,60.261)=1.559, p=0.190). Conversely, the DG (F(13,78)=5.076, p<0.001) and AG (F(13,65)=6.090, p<0.001) groups both significantly decreased and increased responding over days, respectively (Fig. 1D). There were no significant interactions with sex (Fs<1.399, ps>0.165). Responses during extinction tracked the differences seen at the end of self-administration for each group (F(2,21) = 13.012, p<0.001; *Figure 1E*). Bonferroni corrected t-tests highlight significant differences between all groups.

### Taste Reactivity

The number of gapes or licks did not significantly differ between groups (gapes: F(2,17) = 1.611, p = 0.229; *Fig. 2A*; licks: F(2,17)=3.056,p=0.074; *Fig. 2B)*. However, a significant sex x group interaction was found for licks (F(2,17) = 3.589, p = 0.050). DG females (n = 2) had significantly fewer licks than AG females (n = 3), although this finding was based on a very small sample size. Sex differences were not seen in gapes (F(2,17) =1.186,p=0.330). In contrast to gapes or licks alone, there were significant differences in net aversiveness (F(2, 21) = 4.245, p = 0.028). Specifically, Bonferroni post hoc tests demonstrated that DG rats had significantly greater net aversiveness than LG rats (Fig. 2C). There were no sex differences in net aversiveness F(2, 21)=2.102, p=0.147). Finally, there were no significant correlations between the last three days of self-administration (Days 12-14 average) with either licks, gapes, or net aversiveness (rs > - 0.035, p > 0.75). However, the slope of self-administration seen across the 14 days did predict net aversiveness in the DG group. Specifically, animals with the steepest drop in self-administration had the highest net aversiveness (r = 0.79342, p = 0.033; Fig. 2E). Conversely, there was no relationship between net aversiveness and slope for either the LG or AG group (rs < 0.45, ps > 0.17; Fig. 2D and F).

**Figure 2.**
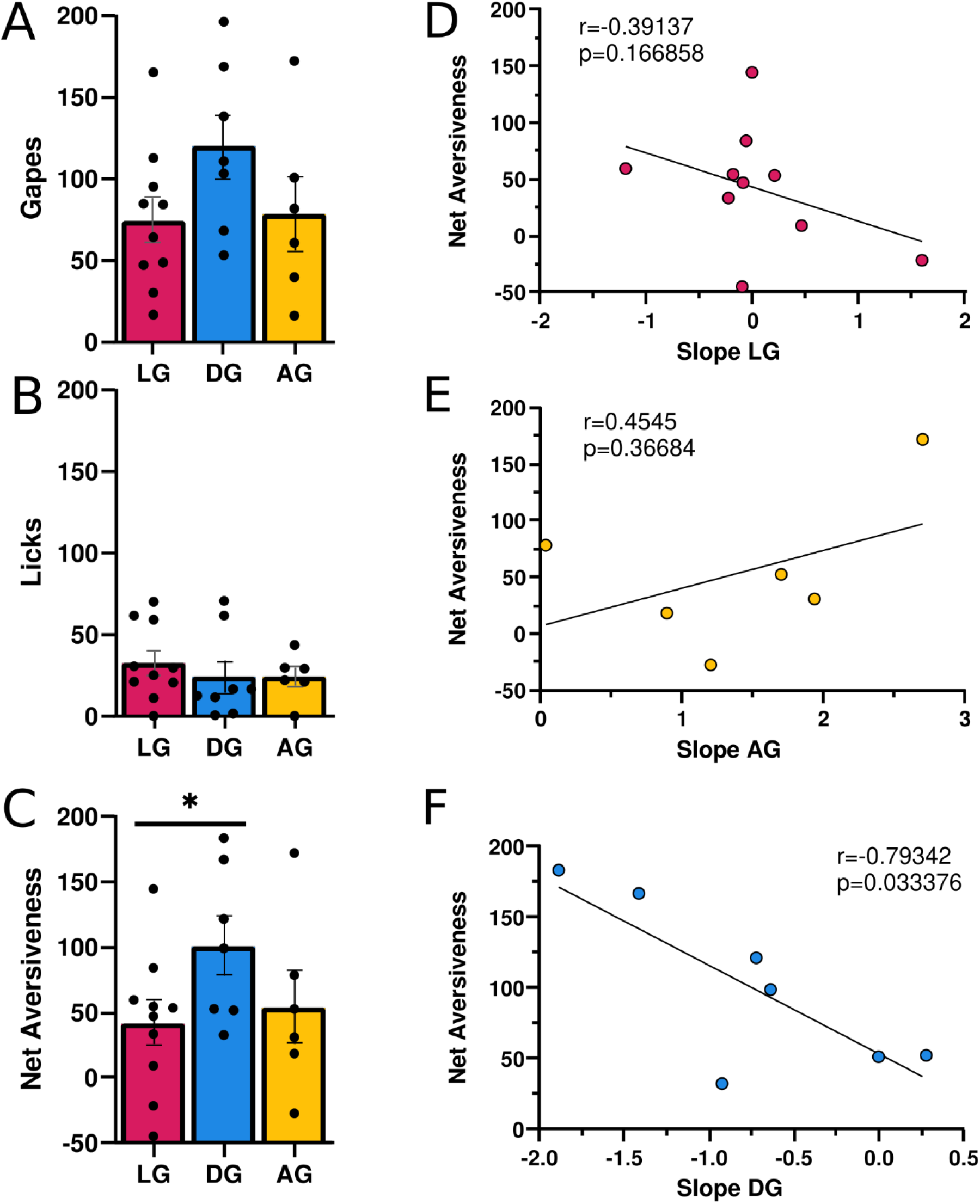
The value of quinine varies depending on self-administration patterns. There was no difference in gapes (**A**) or licks (**B**) for any group. However, DG rats had significantly higher net aversiveness that LG rats (**C**). **D**) There was no relationship between the slope of self-administration behavior and net aversiveness in LG rats. **E**) DG rats with the steepest slope had the highest aversive taste reactivity. **F**) There was no relationship between the slope of self-administration behavior and net aversiveness in AG rats. * p < 0.05

## Discussion

Drug use is often paired with aversive cues. Here, we developed a novel model of cocaine self-administration paired with aversive intraoral quinine. Using this model, we identified two distinct groups of rats that either did not self-administer cocaine or that escalated cocaine intake, similar to divergent patterns of intake seen in human populations. Both groups had similar aversive TR to quinine, suggesting that the escalating group did not habituate to the aversive cue, but rather pursued drug in spite of it. In addition to these two groups, we also identified a third group with high initial intake that subsequently decreased over time. We argue that this third group may represent individuals who “overindulge” on their first use and subsequently find self-administration to be aversive. Together, these data support our new model and provide valuable insight on the impact of aversive cues on self-administration behavior.

Two of the groups we identified generally confirmed our expectations. Quinine appeared to suppress day 1 self-administration in both AG and LG rats (Fig. 1D), especially when compared to the larger amounts normally self-administered on day 1 of cocaine self-administration in tasks using nosepokes (Moschak & Carelli, 2017). However, by Day 14 the AG rats attained similar amounts of cocaine as conventional self-administration. Additionally, the AG rats continued to respond for quinine even during extinction (Fig. 1E). This was in spite of the fact that AG rats exhibited high aversive TR to quinine that was no different than the LG rats (Fig. 2A-C). This suggests that AG rats escalated intake *despite* the aversive quality of quinine. These findings contradict surveys demonstrating that the aversive effects associated with drug intake diminish with repeated use (Friedman et al., 1985; Hahn et al., 1990) and studies reporting that both positive *and* negative experiences are predictors for continued use (Urbán, 2010). However, they support others suggesting that positive experiences alone are predictive (Pomerleau et al., 1998; Wellman et al., 2014). In total, our new task models real-world behavior whereby most individuals do not pursue drugs of abuse (LG rats), while a minority escalate their drug intake (AG rats).

The third group we identified (DG rats) was unexpected. Day 1 cocaine intake for DG rats was higher than previous reports for short-access self-administration (Moschak and Carelli, 2017), then rapidly decreased below typical self-administration by Day 3 (Fig. 1D). The excessive initial intake could have been due to high novelty-seeking for the novel cocaine and quinine sensations, since novelty-seeking has been implicated in heightened acquisition of cocaine intake (Belin et al., 2008). On the other hand, the sharp decrease in intake was likely due to aversion experienced over the first few days. In support of this, the decrease in intake predicted subsequent overall aversive TR in DG rats (Fig. 2F), which was in turn higher than that seen in low intake rats (Fig. 2C). It may be that DG rats’ initial high cocaine intake itself contributed to the increased quinine aversion, as cocaine has delayed dysphoric effects (Ettenberg et al., 1999) that at high doses could have reduced desire for subsequent intake. The existence of an “overindulging” group may have been overlooked in clinical studies, given that human surveys typically report solely on ceased or continued drug use (Miech, 2023; SAMHSA, 2022). However, a similar “excessive use” phenomenon has been used before in “Rapid Smoking” studies as an attempt to decrease nicotine intake (Lichtenstein et al., 1973; Danaher,1977).

In conclusion, our study introduced a novel task which paired aversive quinine with cocaine from day 1 onwards. We believe the three distinct groups we see mimic real-world variability in initial interactions with drugs of abuse. Future research should investigate the neurobiological mechanisms underlying our task, particularly focusing on connectivity between the insula, medial prefrontal cortex, and ventral striatum which are implicated in aversive-resistant motivation and drug/reward-seeking (Peciña and Berridge, 2005; Hurley and Carelli, 2020; Naqvi et al., 2007; Moorman and Aston-Jones, 2015; Moschak et al., 2018; Jing et al.., 2022).

## Acknowledgements

We would like to thank Karla Galvan, Ricardo Sosa, Marina Peveto, and Kimberly Turner for their technical assistance with this study.

## Author contributions

**Rosalie E. Powers:** Data curation, Investigation, Project Administration, Supervision, Writing-original draft, Writing-review and editing; **Peter A. Fogel:** Data Curation, Formal Analysis, Investigation, Visualization, Writing-original draft, Writing-review and editing; **Jayson H. Reeves:** Investigation, Visualization, Writing-original draft; **Pamela Madrid:** Investigation, Visualization, Writing-original draft; **Travis M. Moschak:** Conceptualization, Formal analysis, Funding acquisition, Supervision, Writing-review and editing.

